# The Impact of Task Context on Predicting Finger Movements in a Brain-Machine Interface

**DOI:** 10.1101/2022.08.26.505422

**Authors:** Matthew J. Mender, Samuel R. Nason-Tomaszewski, Hisham Temmar, Joseph T. Costello, Dylan M. Wallace, Matthew S. Willsey, Nishant Ganesh Kumar, Theodore A. Kung, Parag G. Patil, Cynthia A. Chestek

## Abstract

A key factor in the clinical translation of brain-machine interfaces (BMIs) for restoring hand motor function will be their robustness to changes in a task. With functional electrical stimulation (FES) for example, the patient’s own hand will be used to produce a wide range of forces in otherwise similar movements. To investigate the impact of task changes on BMI performance, we trained two rhesus macaques to control a virtual hand with their physical hand while we added springs to each finger group (index or middle-ring-small) or altered their wrist posture. Using simultaneously recorded intracortical neural activity, finger positions, and electromyography, we found that predicting finger kinematics and finger-related muscle activations across contexts led to significant increases in prediction error, especially for muscle activations. However, with respect to online BMI control of the virtual hand, changing either training task context or the hand’s physical context during online control had little effect on online performance. We explain this dichotomy by showing that the structure of neural population activity remained similar in new contexts, which could allow for fast adjustment online. Additionally, we found that neural activity shifted trajectories proportional to the required muscle activation in new contexts, possibly explaining biased kinematic predictions and suggesting a feature that could help predict different magnitude muscle activations while producing similar kinematics.

## INTRODUCTION

Spinal cord injury affects an estimated 296,000 people in the United States (National Spinal Cord Injury Statistical Center, 2021). People with quadriplegia have ranked the restoration of hand and arm function as very important for quality of life (Anderson, 2004; Collinger, Boninger, et al., 2013). Functional electrical stimulation (FES) is a therapy that can restore hand and arm function by electrically stimulating muscles in order to cause contractions. Studies have demonstrated the use of FES to restore at least some hand function since the 1980s (Kilgore et al., 1989; Peckham et al., 1980), which has resulted in commercially available systems such as the Freehand System (Peckham et al., 2001). These systems however typically relied on external motion or myoelectric commands from residual muscles. These control schemes for FES require residual function and can be unintuitive to use, especially when controlling more than one degree-of-freedom.

Brain-computer interfaces (BMIs) have the potential to provide more intuitive control signals for therapies like FES, enabling people with paralysis to interact with a computer or a prosthesis. These BMIs capabilities have been made possible by a history of neuroscience studies finding that motor cortex activity is correlated with a multitude of movement variables, from intrinsic variables like joint angle and muscle activation (Evarts, 1968), to extrinsic movement direction (Georgopoulos et al., 1986). Taking advantage of these correlations allows linear models to predict these movement variables from neural activity. This approach has been used in BMIs to allow non-human primates to control computer cursors (Gilja et al., 2012; Serruya et al., 2002; Taylor et al., 2002), prosthetic arms (Carmena et al., 2003), and FES (Badi et al., 2021; Ethier et al., 2012; Moritz et al., 2008). Additionally, success in animal BMIs led to the use of similar models in clinical trials as well (Ajiboye et al., 2017; Bouton et al., 2016; Collinger, Wodlinger, et al., 2013; Gilja et al., 2015; Wodlinger et al., 2015). These studies, however, are generally performed in a controlled lab environment, and use relatively simple linear models to make predictions. One key factor in the translation of lab BMI FES systems to tasks of daily living will be how robust they are to the varying environment found in patient’s homes. Groups have included object interaction in their tasks, for example grabbing a coffee cup (Ajiboye et al., 2017), or different size objects (Wodlinger et al., 2015), however there has not yet been a systematic effort to understand how task context affects BMI performance.

Offline studies of how motor cortex controls movement have helped to inform how well we can expect BMI models to generalize. This work has shown that the linear encoding of movements in motor cortex can change with many factors such as posture or task duration (Churchland & Shenoy, 2007; Kakei et al., 1999; Naufel et al., 2019; Scott et al., 2001; Sergio et al., 2005). Recent studies have emphasized instead that the role of motor cortex is to generate movements rather than represent movements (Churchland et al., 2012; Russo et al., 2018; Shenoy et al., 2013). In this view, the activations of single neurons are coordinated. The underlying network connectivity constrains population activity to a low-dimensional manifold and activations on this low-dimensional manifold then form the basis for neural dynamics which generate movements (Gallego et al., 2017; Shenoy et al., 2013). A key feature of these dynamics that is different from a representation model is that they may have a more computational function, for example ensuring that outgoing commands can be generated reliably (Russo et al., 2018). The resulting activity may then change when the same movements are done in different ways because a different computation is needed to generate the movements. As a result, an individual neuron’s activity, which is related to the latent activity in this low-dimensional manifold, could correlate with movements differently when the task is changed to one that requires different neural dynamics. With respect to BMI applications, the decoding models assuming a linear relationship, and non-linear models that do not account for these changes, would then be unable to make accurate predictions in the new tasks.

It is still unclear how large of a task change will require different neural dynamics and thus a different decoding strategy. It has been shown that different dynamics are required for large changes in a task, such as forward versus backward arm pedaling (Russo et al., 2018), reaching or walking (Miri et al., 2017), or using one arm or the other (Ames & Churchland, 2019). At the same time, there is evidence that tasks with the same movements performed differently may have similar neural dynamics. A recent study found that cycling at different speeds led to similar elliptical trajectories in high variance neural dimensions, with a lower variance dimension encoding task speed (Saxena et al., 2022). Additionally, a study of isometric, resisted, and free-moving wrist movements found a neural manifold that explained a large amount of neural variance in all tasks (Gallego et al., 2018) and a similar study comparing the same wrist movements found that they could still predict muscle activations between the contexts although it required a gain-factor related to required muscle activation (Naufel et al., 2019). These observations suggest that with small variations to a task, such as a change in speed or muscle exertion, the neural dynamics may be similar across task variants, with variation in lower variance dimensions.

Which tasks require a change in neural dynamics is a particularly important question to study for hand movements, as the hand is the major end effector interacting with the environment in varying postures and with different loads. However, this work has not yet been extended to continuous finger movements. Finger movements are less studied than arm reaches but initial studies show that grasping movements may show different dynamics due to the increased proprioceptive and tactile feedback present (Goodman et al., 2019; Suresh et al., 2020). In a promising start to studying decoder generalization for individuated finger movements, it has been shown previously that multiple finger movements can be predicted simultaneously, in real-time, and that a linear model trained with data from individual finger movements could also predict combined finger movements (Nason et al., 2021), suggesting that individual finger movements and combined finger movements may have similar neural dynamics.

In this study, we investigate how well the decoding of finger movements from intracortical neural activity in nonhuman primates can generalize to realistic alterations of the context in which a task is performed, similar to those that may be found in a BMI user’s home. We ask how the relationship between intracortical neural activity, non-prehensile finger movements, and the related muscle activations are impacted by context changes, such as spring-like resistances and postural changes. We show that these context changes reduce our ability to predict finger kinematics and finger-related muscle activations offline. However, in an online kinematic-based finger BMI task, the monkey can accommodate for the changed task context and achieve near equivalent performance with or without the context change. We explain this by showing that the underlying neural manifolds stay well aligned between contexts, the neural dynamics are shifted due to context, and the shift in neural dynamics can be related to the muscle activation required in the new context.

## RESULTS

### Context changes alter muscle activations and neural activity

We are ultimately interested in understanding the impact of context changes, such as wrist flexion or spring resistance, on BMI decoding performance. We first asked whether introducing context changes during a controlled task causes any change in behavior, muscle activation, or neural channel activation. Our first manipulations were the addition of torsional springs or the static flexion of the wrist by 23 degrees, referred to as the spring and wrist contexts respectively, during a 1-degree-of-freedom (1-DOF, all fingers flex or extend together) center-out task. The torsional springs resist flexion such that more force is required to flex the fingers but less force is required to extend the fingers. We expected the springs to cause minimal to no change in finger velocity during movements but a large increase in muscle activation for flexor muscles during flexion as well as a decrease in activation for extensor muscles during extension. Figure 1A shows finger position and velocity traces averaged over all flexion trials (solid lines) and all extension trials (dashed lines) on one representative day for monkey N where the spring manipulation was tested. We see small changes between the velocities in normal trials (black traces) and spring trials (blue traces). To quantify this change, we compared the peak velocities between normal trials and other context trials for representative sessions with each context (Figure 1C). We found that the largest changes in peak velocity were monkey N extending fingers 12.5% faster during wrist trials (p = 5e-24, two-sample t-test), and monkey W flexing fingers 22.3% faster during wrist trials (p = 2.6e-11, two-sample t-test), with both monkeys showing small changes in peak movement velocity for at least one movement in each context (p<0.05, two-sample t-test).

**Figure 1.**
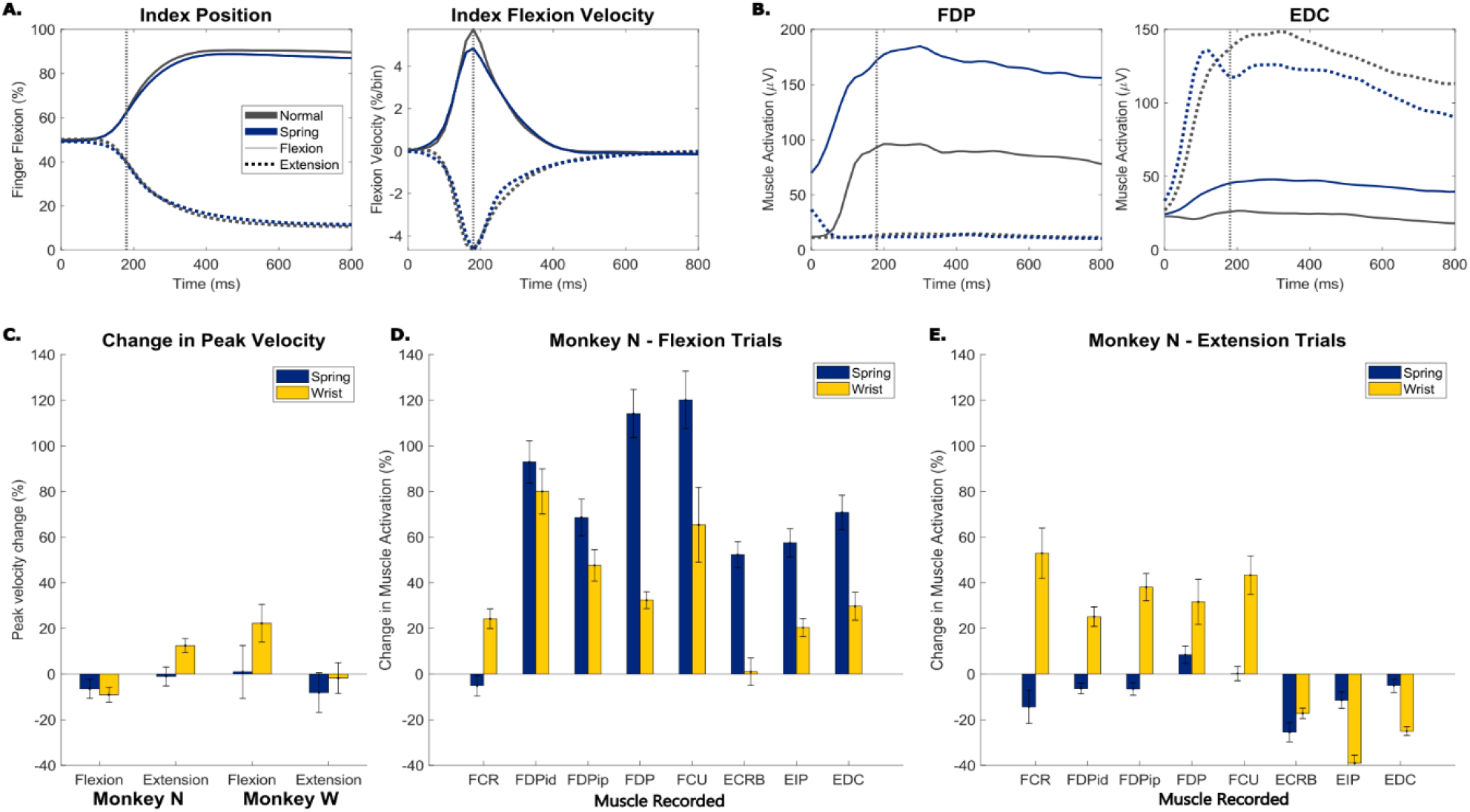
(A) Trial-averaged traces of index finger position and index finger flexion velocity for an example spring session with monkey N. Trials are aligned to peak movement (vertical grey line). Black traces are normal trials, blue are trials with springs in the manipulandum, solid traces are for flexion trials, and dotted traces are for extension trials. (B) Trial-averaged traces of flexor digitorum profundus (FDP) muscle activation and extensor digitorum communis (EDC) muscle activation for an example spring session with monkey N. Formatted the same as (A). (C) Change in peak velocity between normal trials and trials with either springs present or the wrist flexed for both monkey N (left) and monkey W (right). Trials are split by movement direction, either flexion or extension. Error bars indicate 99% confidence interval based on a two-sample t-test. (D, E) Change in average muscle activation in a window around peak movement between normal trials and trials in the spring (blue) or wrist context (yellow) for all eight muscles recorded in monkey N. Trials are split between flexion (D) and extension (E) movements. Error bars indicate 99% confidence interval based on a two-sample t-test.

In contrast, during the same 1-DOF task, muscle activations change substantially for trials toward both targets (Figure 1B), showing trends such as increased flexor digitorum profundus (FDP) muscle activation for flexion and less extensor digitorum communis (EDC) activation for extension. All muscles implanted are included in Table II (Methods). We compared the average muscle activation in a window around peak movement between normal trials and off-context trials for the same representative sessions for Monkey N (Figures 1D, 1E). During spring trials, we found that every muscle except FCR required significantly higher than normal trial muscle activation for flexion (Figure 1D blue, p<0.004 two-sample t-test), an average increase of 91.9% for the finger flexor muscles (FDPid, FDPip, FDP), and every muscle except FDP and FCU required less muscle activation for extension (Figure 1E blue, p<1e-5 two-sample t-test), an average decrease of 8.2% in finger extensor muscles (EIP, EDC). During wrist trials, finger flexor muscles showed an average 53.3% increase in activation for flexion trials (Figure 1D yellow) and finger extensor muscles had an average 32% decrease in activation for extension trials (Figure 1E yellow).

After establishing that context had a large effect on muscle activity with a relatively small effect on finger kinematics, we next evaluated whether neural activity changed due to the addition of springs or altered wrist posture. For each neural channel we recorded two features, the threshold crossing firing rate (TCFR), and spiking band power (SBP). SBP is a low power feature that has been previously shown to be well correlated with the firing rate of the largest amplitude unit (Nason et al., 2020), often enabling us to identify more tuned channels. We evaluated how many channels were tuned to movement and how many of these tuned channels modulated activity with context change as described in methods. The results are included in Table I. The SBP feature resulted in an average of 86.9 and 28 tuned channels of 96 for monkey N and monkey W respectively, while TCFR resulted in an average of 36.7 and 11.8 tuned channels of 96 for monkey N and monkey W respectively. An average of 24.4% of the tuned TCFR channels and 37.7% of the tuned SBP channels significantly changed activity with the wrist context and an average of 56.8% of the tuned TCFR channels and 52.3% of the tuned SBP channels significantly changed activity with the spring context. As both features had a similar proportion of tuned channels that were modulated by context changes, we opted to use SBP as the primary feature for the subsequent analyses in order to increase the number of tuned channels available for analysis.

**Table I.** Number of channels tuned to any movement using two features (TCFR and SBP) for four types of experimental sessions (Monkey N or Monkey W with the spring or wrist context) and the percentage of tuned channels that showed a significant change in trial feature during either flexion or extension trials of the other context on that experimental session compared to normal trials. Standard deviation (SD) is calculated across sessions of the same type and n is the number of sessions of that type which is indicated in the first column.

|  | TCFR |  | SBP |  |
| --- | --- | --- | --- | --- |
|  | Tuned Channels | % Context Modulated Channels | Tuned Channels | % Context Modulated Channels |
| Monkey N Spring Days (n=3) | **38.3** (SD=4.0) | **47.0%** (SD=2.8) | **89.3** (SD=2.1) | **61.7%** (SD=12.0) |
| Monkey N Wrist Days (n=2) | **35.0** (SD=1.4) | **43.1%** (SD=9.8) | **84.5** (SD=7.8) | **54.4%** (SD=0.8) |
| Monkey W Spring Days (n=3) | **12.0** (SD=1.0) | **66.5%** (SD=2.8) | **35.0** (SD=6.9) | **42.8%** (SD=7.1) |
| Monkey W Wrist Days (n=2) | **11.5** (SD=3.5) | **5.6%** (SD=7.9) | **21.0** (SD=5.7) | **20.9%** (SD=4.5) |

### Decoding neural activity across task context

After confirming that these context changes had large impacts on muscle activation (Figure 1) and affected many channels of neural activity (Table I), we next asked how this will impact the ability to decode intended movements for BMI applications. Typically, BMIs use linear models to relate neural firing rates to the desired control variable (Ajiboye et al., 2017; Nason et al., 2021; Wodlinger et al., 2015). Given the work showing that task changes similar to those tested here can alter the linear encoding of muscle activations (Naufel et al., 2019) in motor cortex, we next ask if the same is true for individuated finger movements. To test this, we recorded kinematics for both monkeys and muscle activations for monkey N during a two-finger task and then trained linear models with data from normal trials to predict muscle activations or kinematics in unseen normal trials or other context trials.

We first present the results for decoding muscle activations across context. Figure 2A shows average predictions of FDP and EDC muscle activations for normal trials and spring context trials from one example experimental day, both using a linear model trained on normal trials. We found that within-context linear models, i.e. models trained and tested on the same context trials, could predict muscle activations well during individuated finger movements, with average correlations comparable to predictions of kinematics (supplementary Tables I, III). Models trained on normal trials however are consistently unable to predict muscle activations well in the off-context trials. For example, when springs are present the predictions do not account for the large changes in FDP activation magnitude or EDC activation during flexion trials (Figure 2A). Across three sessions in each context – wrist, spring, or both wrist and spring – prediction mean-squared error (MSE) increased significantly from the normal trial baseline both for flexor muscles (FDP, FDPip) and extensor muscles (EDC, EIP) (evaluated by paired t-test, p<2e-9), in each tested context, with an average increase of 188.7% across all context changes and muscles. The increases in error varied widely, ranging from 21% increase (flexors with wrist-flexed) to 356% increase (flexors with both wrist-flexed and springs).

**Figure 2.**
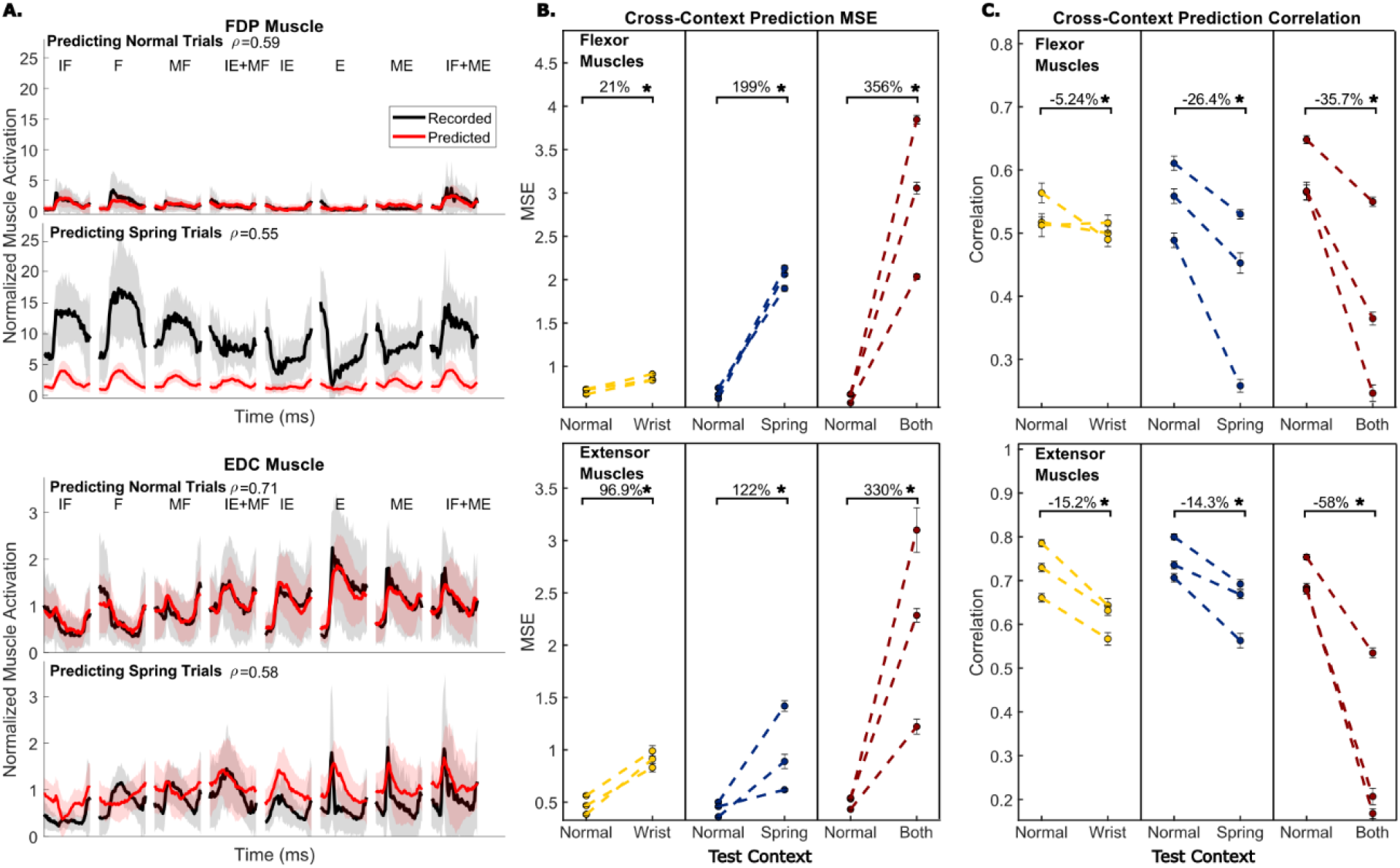
(A) Recorded and predicted muscle activation traces for FDP muscle (top half) and EDC muscle (bottom half) from one example session. Traces are aligned to peak movement and averaged over trials to the same trial, shading represents one standard deviation. Predictions are from a model trained only on normal trials and the model is evaluated either on normal trials (top) or trials with springs present (bottom). IF – index flexion, MF – MRS flexion, F – both fingers flexion, IE – index extension, ME – MRS extension, E – both fingers extension. (B) Change in prediction MSE when a model trained on normal trials is evaluated on trials in a different context. Color indicates which context is being tested, yellow is wrist, blue is spring, and both is red. The dashed lines connecting dots represent that the same model was used for both measurements. Error bars on the dots indicate one standard deviation for model performance calculated with 10-fold cross-validation. (C) Same as (B) but model performance is measured with prediction correlation.

We next asked whether this dramatic increase in prediction error is driven by a simple offset or magnitude change, or a reduced linear relationship with recorded muscle activations. For example, the off-context predictions of FDP activation during flexion in Figure 2A do not account for the offset in muscle activation at the beginning of trials and do not predict a large enough change in the magnitude of muscle activation throughout the trial, but the same linear correlation is maintained. Alternatively, the off-context predictions of EDC activation during flexion are less correlated with measured EDC activation because the model predicts EDC inactivation, as is the case during normal trials, even though spring trial EDC activation increases for flexion. Since MSE is influenced by both factors, we also tracked prediction correlation (Figure 2C) which is less affected by offsets and magnitude differences. When we evaluated model performance with correlation, using the same sessions and models trained on normal trials as when measuring MSE, we found that prediction correlation decreased from normal baseline by an average of 25.8% across tested contexts and muscles. This change was significant for all tested contexts (paired t-test), ranging from a 2.4% decrease for FDPip in the wrist context (p=0.03) to a 69.4% decrease for EDC in the both context (p=1.3e-17). While significant, the change in correlation was a smaller effect than the change in MSE.

Kinematics are used as a control signal in BMI applications more frequently than muscle activation so we next examined the error in predicting finger position and velocity across contexts. Figure 3A shows trial-averaged predictions for each target from training a linear model on normal trials and predicting normal trials or spring trials for an example session with monkey N. In both index and MRS flexion predictions, we observed the off-context predictions to be worse than the normal trial predictions. Predictions during the spring trials often showed a bias towards flexion. We measured changes in prediction accuracies on three days for each context – spring, wrist, and both – for monkey N, and one additional day for each context for monkey W (Figures 3B and 3C). All context changes resulted in significantly higher prediction MSE (paired t-test, p<1e-4), averaging 68.2% for finger position and 11.4% for finger velocity. All context changes also resulted in small but significant decreases in prediction correlation (paired t-test, p<1e-4), averaging −18.6% for finger position and −12.8% for finger velocity. The smaller change in prediction correlation indicates that much of the prediction error is coming from offsets or magnitude differences in the predictions.

**Figure 3.**
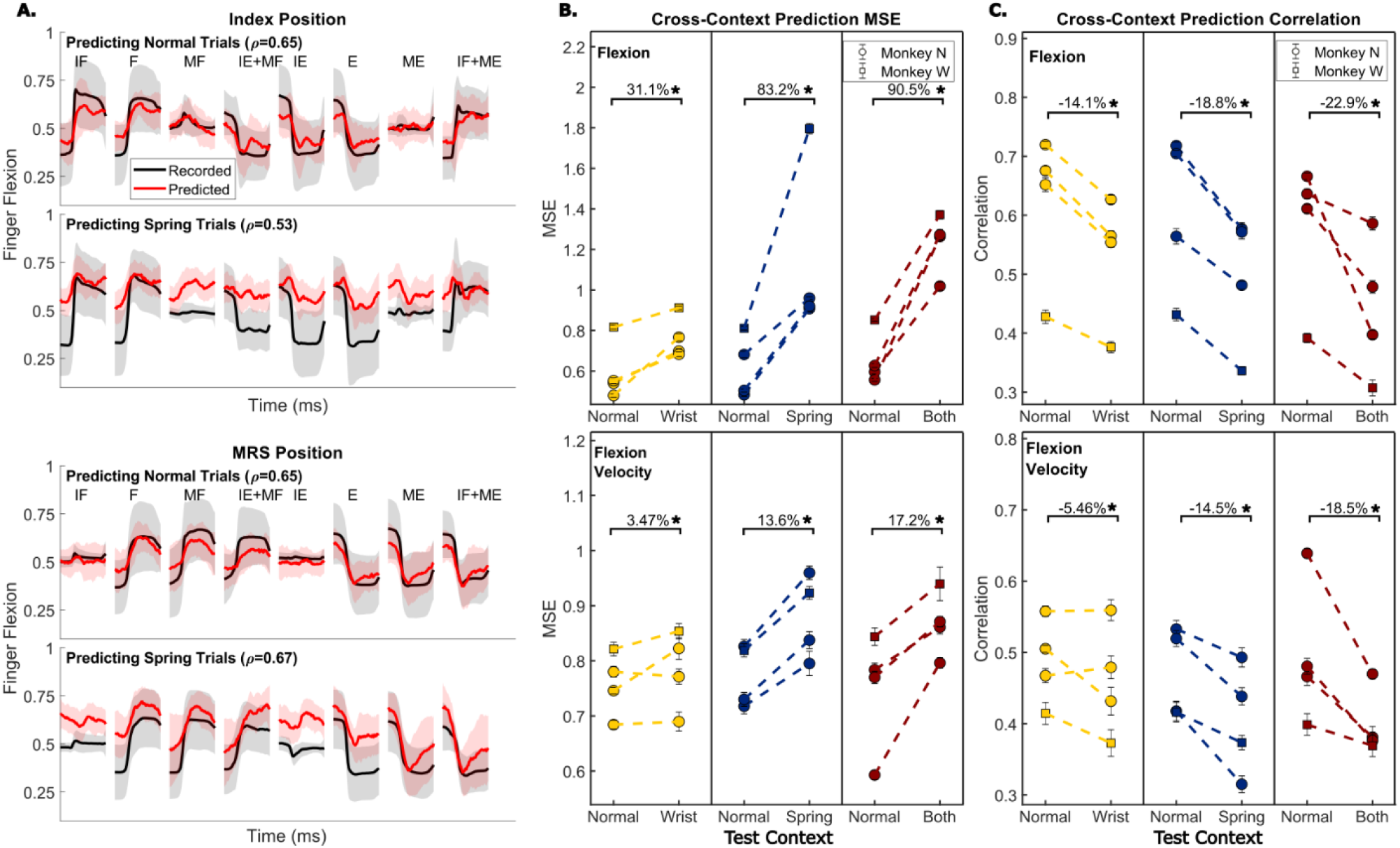
(A) Recorded and predicted position traces for index finger position (top) and MRS finger group position (bottom), averaged across all trials towards each target. Shading represents one standard deviation. Predictions are from a model trained only on normal trials and the model is evaluated either on normal trials or trials with springs present. (B) Change in prediction MSE when a model trained on normal trials is evaluated on trials in a different context. Color indicates which context is being tested, yellow is wrist, blue is spring, and both is red. The dashed lines connecting dots represent that the same model was used for both measurements. Error bars on the dots indicate one standard deviation for model performance calculated with 10-fold cross-validation. (C) Same as (B) but model performance is measured with prediction correlation.

### Changing task context has minimal impact on online BMI performance

Based on these offline prediction results, we might expect that in a real-time BMI, a model trained on normal trials will be more difficult to use when controlling a virtual hand in a new context. We investigated this by training either a Kalman Filter (KF) or a ReFIT Kalman Filter (RFKF), as done previously by Nason et al. (2021), and having the monkey control the virtual hand with the model while we applied context changes to this virtual task. We introduced context changes in two separate ways. First, we added springs, a static wrist flexion, or both to the manipulandum and had the monkey control the virtual hand with a RFKF trained on normal trials. Second, we trained different KFs using training data collected in different contexts and had the monkeys use the KFs in the online task without any context changes applied to the manipulandum. Due to the quality of recorded neural signals, Monkey W controlled only 1-DOF online while Monkey N controlled 2-DOF online.

We first tested whether online BMI performance changed when using a standard RFKF with context changes added to the manipulandum, i.e. manipulating the hand context while only neural activity was driving the virtual hand. One RFKF model could be tested on multiple context changes in a single session, for example, Figure 4A shows the acquisition times during an experimental session where two contexts, spring and wrist, were tested. Figure 4B summarizes the changes in online performance over six experimental sessions for monkey N and four sessions for monkey W. During these 10 sessions the context changes were tested 15 times. Each bar compares the performance between normal trials and one off-context condition in a session when using the same model for both. Ultimately, both monkeys were able to consistently reach the same levels of performance despite added context changes to the manipulandum. Of the 15 tests, only one test resulted in a significant change in at least one of the performance metrics (p<0.01, two-sample t-test). In this case, monkey N using the RFKF while his wrist was flexed resulted in a 13.0% increase in time to target (p=6.7e-3), the equivalent of 86ms. This overall lack of change was surprising since the offline decoding results had greater prediction error.

**Figure 4.**
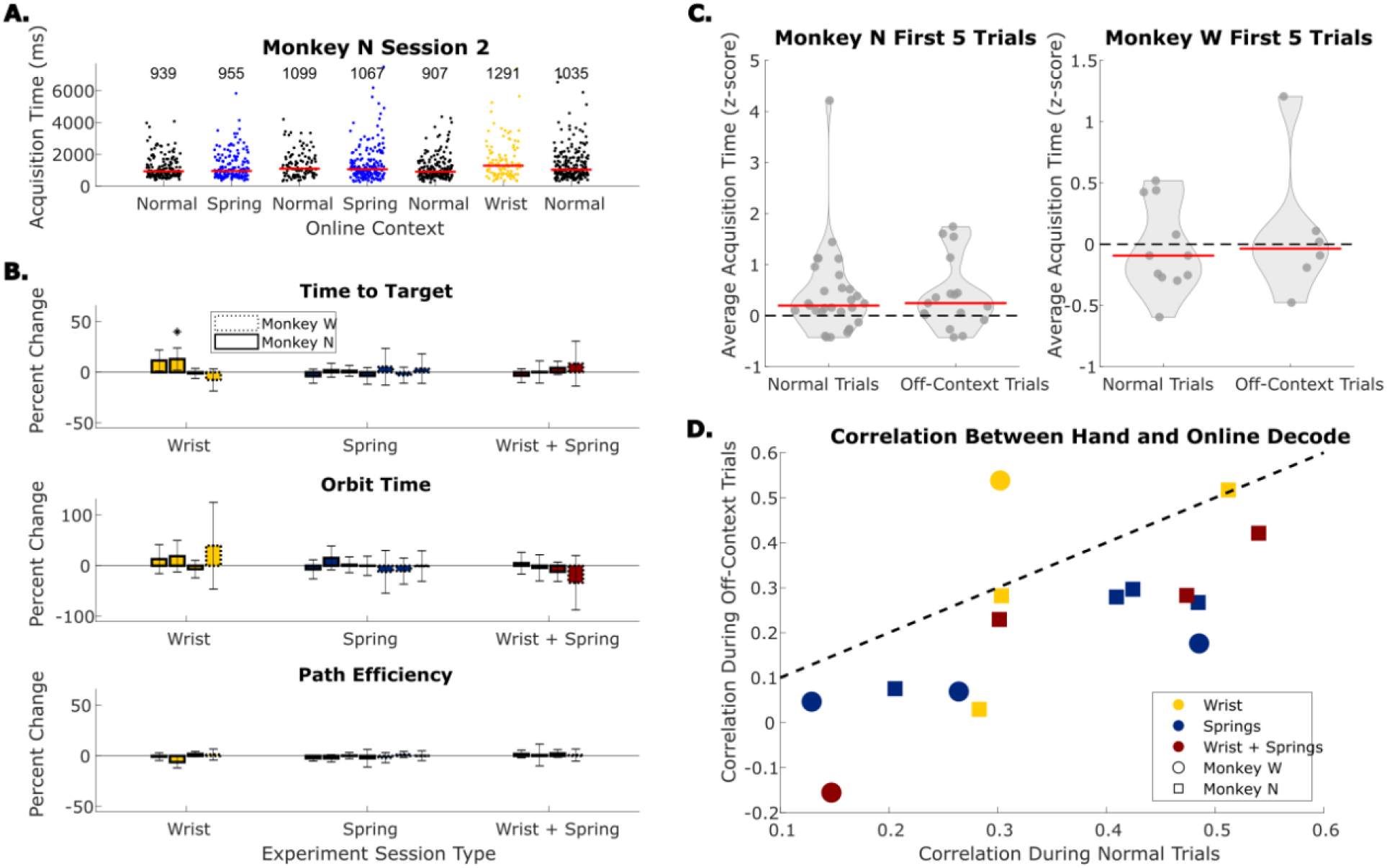
(A) Example online session in which both the spring and the wrist context are tested. Each dot indicates acquisition time of one trial, each grouping of dots is a series of trials before the context was changed. Red bars and numbers above each grouping illustrate the median acquisition time (in ms) for that series of trials. (B) Change in performance metrics between normal online trials and online trials with the context change indicated by the x-axis in the manipulandum. Off-context online trials are compared to the normal online trials immediately before and after them. Error bars indicate 99% confidence interval in performance metric change. (C) Average acquisition time during the first 5 trials each time online trials were started, split between normal trials and trials with context changes applied to the hand (off-context). Acquisition times were z-scored within a series of trials in the same context. Red lines indicate the median. (D) The correlation between hand position and online decode during the trials for each online comparison in (A). Colors represent the type of context change, matching those in (B). Dotted line indicates equal hand-decode correlation in both types of trial.

To explain this minimal change in online performance, we examined each monkey’s ability to adapt to context changes during the online task. Figure 4D shows the correlation between real finger movements and the virtual finger movements during online control during trials with context changes (y-axis) or normal trials (x-axis) for each of the 15 tests in Figure 4B. The monkey’s finger movements were consistently more correlated with the virtual hand during normal online trials. This indicates that during the off-context online trials they adjusted and moved their hand differently to account for the context change. Additionally, to measure the ease of adjustment, we compared the average acquisition times for the first five trials after the start of online trials. Figure 4C shows the distribution of the average acquisition time during the first five trials every time the online trials were started, split between normal trials and off-context trials. Monkey N had a short adaptation period as the average acquisition time during the first five trials was significantly greater than zero (p=0.002, one-sample Kolmogorov-Smirnov test). Monkey W, on the other hand, did not have significant adaptation during the first five BMI trials (p=0.22, one-sample Kolmogorov-Smirnov test). Interestingly, the performance in the first five off-context trials is not different from normal trials for both monkeys (two-sample Kolmogorov-Smirnov test, p=0.88 for monkey N, p=0.79 for monkey W). This suggests that adaptation to a context change is a similar difficulty as adaptation from hand control to BMI control without any context change.

To better isolate the effects of context changes on our online task, in a second online experiment, the monkeys alternated between using two Kalman filters: one trained on normal trials and another trained on off-context trials. Monkeys N and W performed these tests on nine and six separate days respectively. On each day, only two models were trained in order to compare one context change. Figure 5A shows an example session alternating between a model trained on normal trials and a model trained on wrist trials. In 15 sessions, this experiment revealed small but significant changes in at least one performance metric for 11 sessions (Figure 5B, p<0.01, two-sample t-test) although only two sessions had worse online performance for all three metrics. The significant decreases in performance averaged 32.6% for time to target, 46.5% for orbit time, and 8.5% for path efficiency, with the “both” context having the largest effect. As the off-context online performance was worse in the two-model tests, we next asked if this could have been caused by the monkeys not being able to adapt in the same ways as the online context change experiment. Using the same correlation metric between measured finger positions and decoded finger positions as Figure 4D, we found that both monkeys still moved their hand differently between normal trials and off-context trials, although the change was smaller for monkey N. Monkey W actually moved his hand along with the virtual hand more during off-context trials. Finally, we compared the acquisition time in the first five trials while using the normal model or an off-context model (Figure 5C). Similar to Figure 4C, monkey N had a short adaptation period in the first five trials (p=0.008, one-sample Kolmogorov-Smirnov test) that is the same between using the normal model and off-context models (p=0.99, two-sample Kolmogorov-Smirnov test). Monkey W once again did not show a significant adaptation period (p=0.17, one-sample Kolmogorov-Smirnov test) which was the same between using the normal model and off-context models (p=0.5, two-sample Kolmogorov-Smirnov test).

**Figure 5.**
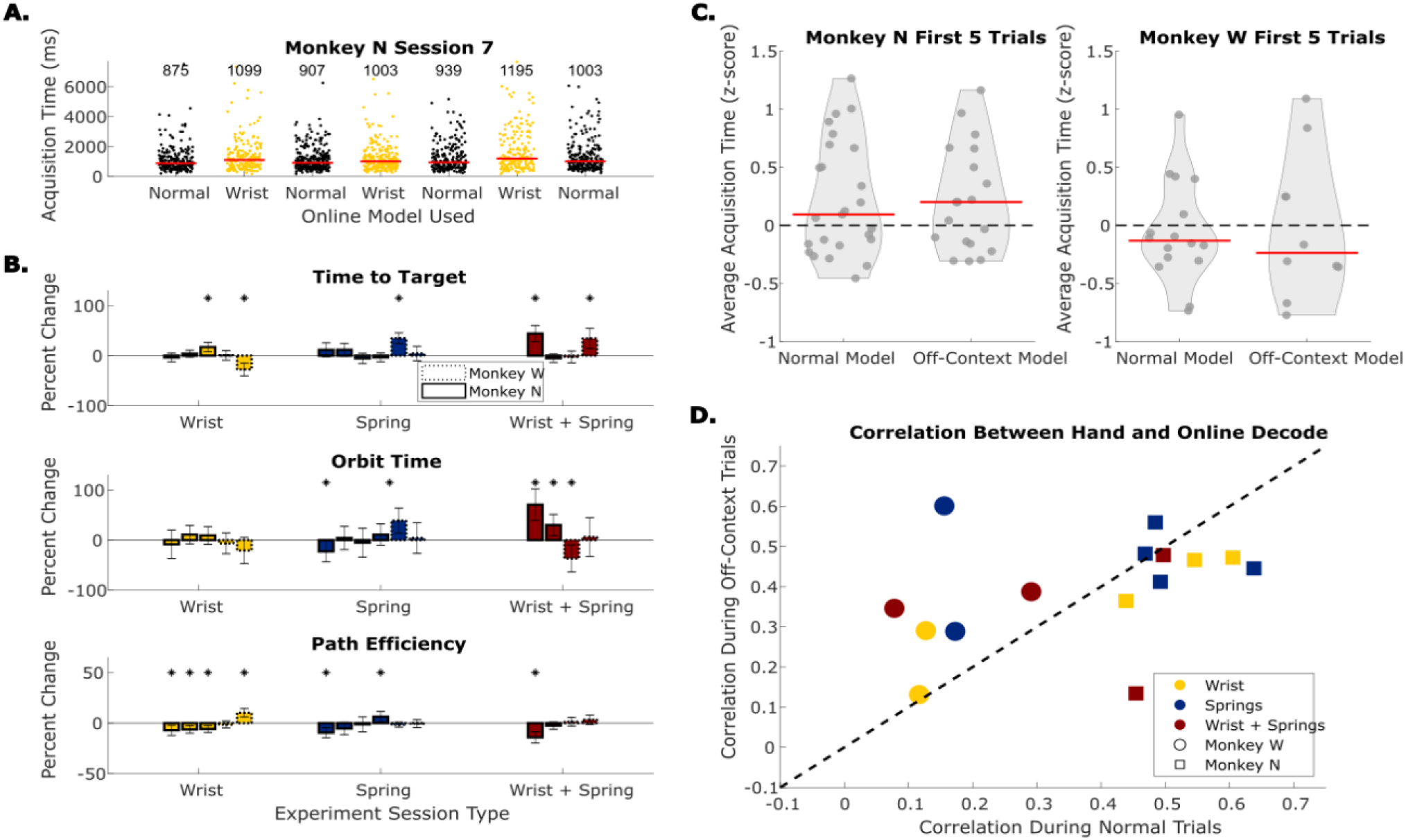
Example online session in which the wrist context are tested. Each dot indicates acquisition time of one trial, each grouping of dots is a series of trials before the context was changed. Red bars and numbers above each grouping illustrate the median acquisition time (in ms) for that series of trials (B) Change in performance metrics between normal online trials and online trials with the context change indicated by the x-axis in the manipulandum. Off-context online trials are compared to the normal online trials immediately before and after them. Error bars indicate 99% confidence interval in performance metric change. (C) Average acquisition time during the first 5 trials each time online trials were started, split between trials performed with the normally trained model and the model trained on off-context trials (off-context). Acquisition times were z-scored within a series of trials with the same model. Red lines indicate the median. (D) The correlation between hand position and online decode during the trials for each online comparison in (A). Colors represent the type of context change, matching those in (B). Dotted line indicates equal hand-decode correlation in both types of trial.

### Context Shifts Population Neural Activity

To help explain how the monkeys were able to adjust to different contexts so well during the online task, we further examined changes in neural activity during the offline task in different contexts. First, we ask if there are any obvious trends in how the channel activity changes during simple 1-DOF movements, for example increasing neural activation when flexion requires more muscle activation. In one experimental session, monkey N performed the 1-D task normally as well as in the wrist, spring, and rubber band contexts. The rubber bands altered the required muscle activations for the task in the same way as the springs, however to a larger extent. In two additional sessions, Monkey W performed the 1-D task normally as well as in the wrist context in one session and spring context in the other session. Figure 6A shows trial-averaged neural activation traces from two example modulated channels, one from each monkey, both comparing the activation during spring trials and normal trials. We found that neural channels showed a mix of changes with context. For example, Monkey W’s channel 12 was activated more compared to normal for spring flexion targets (blue solid), similar to the muscle activations of the finger flexors. However, other example channels like Monkey N’s channel 90 show less neural activation during movement in the spring contexts for both flexion and extension targets which is not consistent with any changes in muscle activation or kinematics.

**Figure 6.**
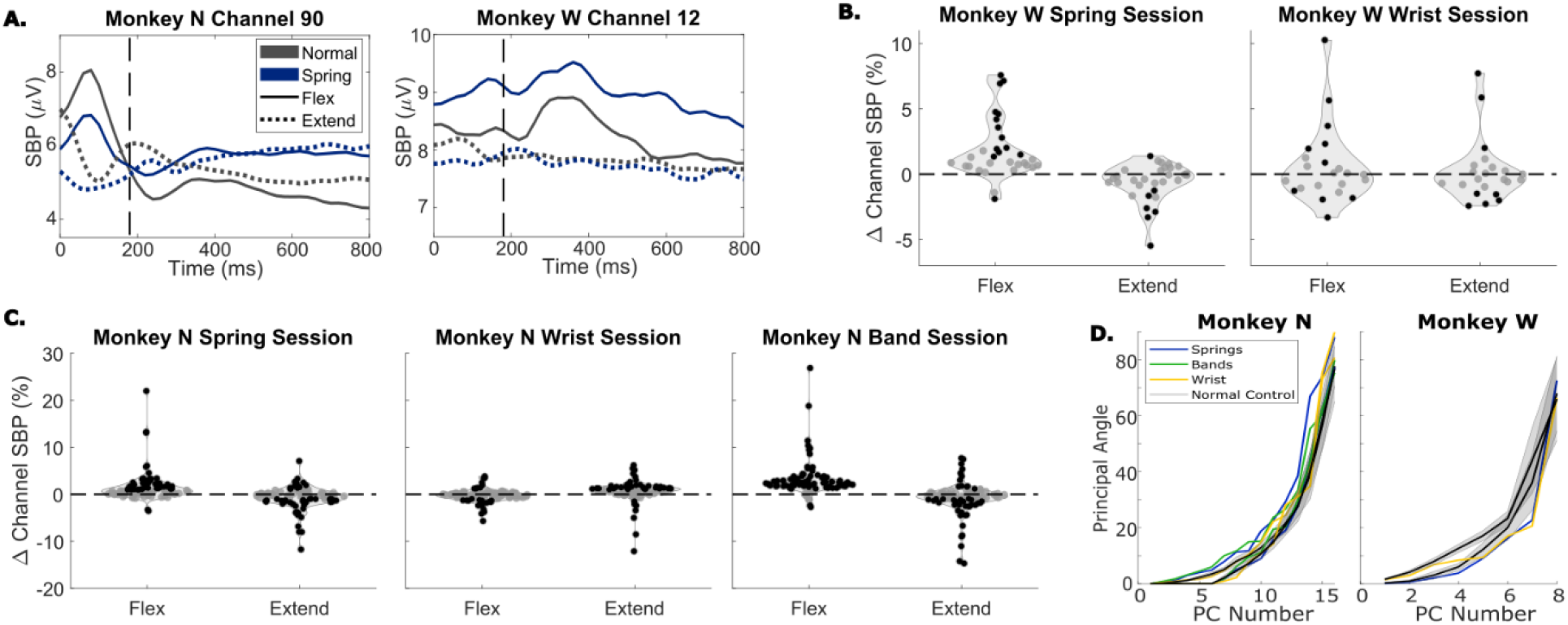
(A) Trial-average SBP for two channels during a spring session for each monkey. Trials were aligned to peak movement before averaging, vertical dashed lines indicate peak movement. Black traces show normal trials, red traces show trials with springs present, solid traces are flexion trials, and dotted traces are extension trials. (B) Change in average SBP in a window around peak movement for each tuned channel for a spring session (left) and wrist session (right), both for monkey W. Black dots indicate significant differences according to a two-sample t-test (p<0.01). (C) Same as (B) but for monkey N. (D) Principal angles between the PCA space calculated for normal trials and the PCA spaces calculated for spring trials (blue), band trials (green), or wrist trials (yellow). Black lines are average angles between PCA spaces for two random sets of normal trials, with grey shading indicating one standard deviation. Solid lines are for session 1 and dashed are for session 2.

To quantify changes in activation for the population of tuned channels, we compared the average channel activation in a window around peak movement for each type of trial. Figures 6B and 6C show the change in SBP for all tuned channels between off-context trials and normal trials, split by flexion and extension trials, for monkey W and monkey N respectively. Black dots indicate channels with significantly different trial SBP between off-context and normal trials towards that target according to a two-sample t-test (p<0.01). During spring trials, context modulated channels were activated significantly less on average for extension and were activated significantly more for flexion with both monkeys (p<0.01, paired t-test). During wrist trials, context modulated channels were activated significantly less for extension for monkey N only (p<0.01, paired t-test) and there were no trends for channels increasing or decreasing activation on average for flexion or extension for monkey W. Notably, the majority of changes in neural activation are on the order of 10% or less for individual channels, fairly small relative to the large changes in muscle activation observed in Figure 1, which is consistent with the results from Naufel et al. (2019) with wrist movements.

We next investigated how consistent the covariance structure of the neural activity is across different task contexts. We calculated the principal components (PCs) underlying the neural activity in each context in order to obtain one manifold for each context. We then found the minimum angles, also known as the principal angles, required to align the PCs from each type of off-context trial with the PCs calculated from normal trials (Figure 6D), similar to what has been previously presented (Gallego et al., 2018). Principal angles were also calculated between manifolds calculated from random sets of normal trials to create a set of control angles. Two sessions with normal, spring trials, wrist flexed, and rubber band trials in the same session are included for monkey N (Figure 6D Left), and two sessions, one with normal and wrist flexed and one with normal and spring trials, are included for monkey W (Figure 6D Right). We found that the principal angles between off-context trial neural activity and normal trial neural activity match the normal control angles well. This indicates that the activity in each context falls within well-aligned manifolds.

Next we looked at how much variance in neural activity is due to the context changes. We calculated 16-dimensional demixed PCA (dPCA, Kobak et al., 2016) components for a neural manifold spanning neural activity during trials from all contexts in a single session for all the days included in Figure 6D. Figure 7A shows the dPCA components for one session with monkey N. The components are organized in rows according to which behavioral parameter they explain the most variance for: time (condition-independent), context, target, or context-target interaction. The amount of variance explained by each behavioral parameter is summarized in Figure 7B for both monkeys. The condition-independent and target parameters together explain the majority of the neural variance. On average, target explains 37.72% of neural variance, the condition-independent parameter explains 47.5% of neural variance, and the context and context-target interaction parameters together explain 24.1% of neural variance for monkey N and 5.5% of neural variance for monkey W. In inspecting the activation of components in Figure 7A, the context related components add a shift to neural activity before and after movement (component #3) and separate normal and wrist trials from spring and rubber band trials during movement (component #6). This creates the picture of dynamics that are largely the same between context but slightly shifted, perhaps in response to a change in proprioceptive input or to generate more muscle activation. We compared the average activation around peak movement of the context-dependent dPCA component that explained the most neural variance with the average muscle activation of flexor or extensor muscles during flexion or extension trials in each context. Figure 7C shows this comparison for three sessions with monkey N, note one session only included normal, spring, and rubber band trials (no wrist). Interestingly, we found that the first context-dependent component correlated very strongly with the activation of the active muscle (i.e. flexor muscles during flexion or extensor muscles during extension) across contexts, with correlations of 0.91 for flexor muscles and −0.89 for extensor muscles. This result may suggest that a feature from neural activity could be used to account for changes in required muscle activation between contexts.

**Figure 7.**
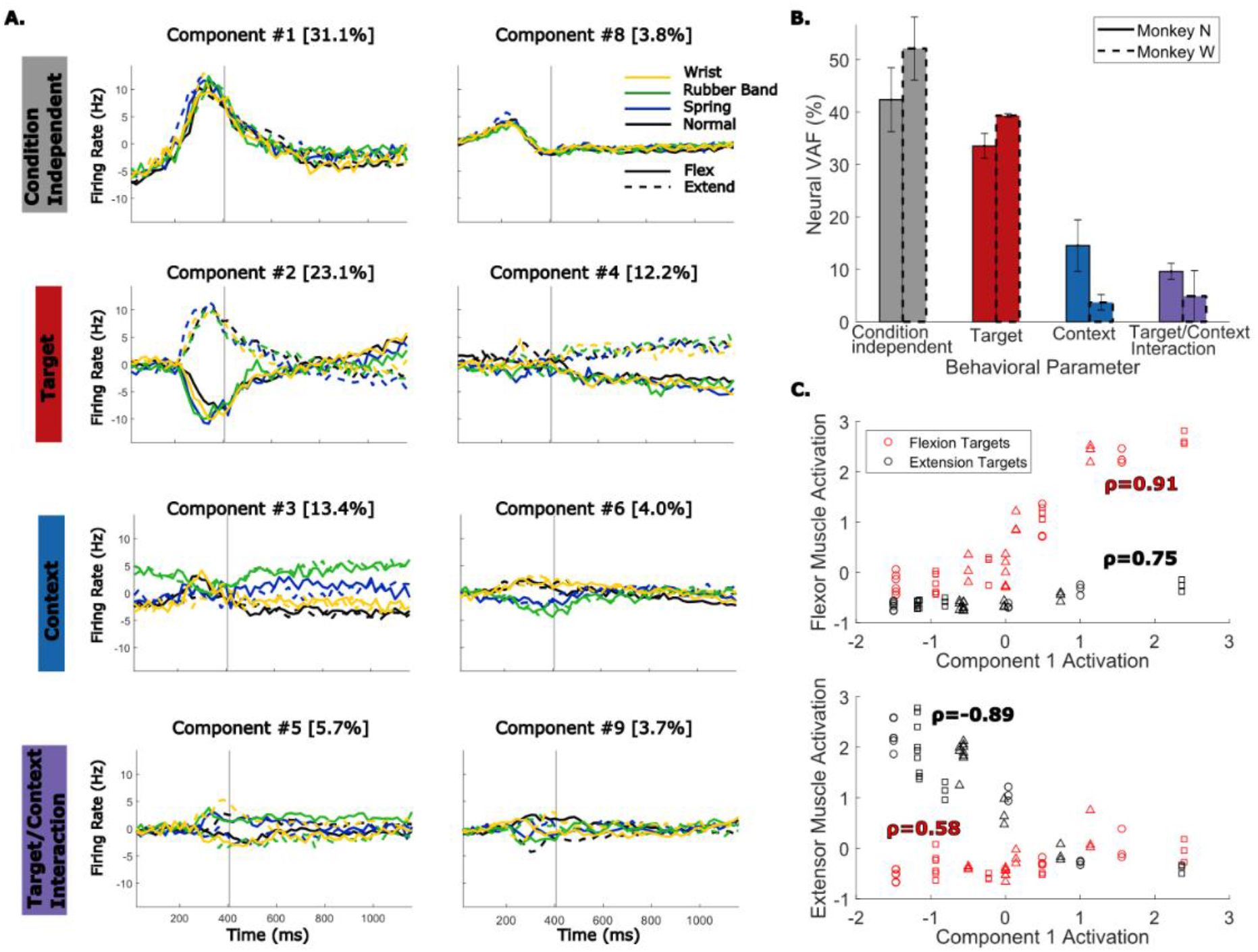
(A) Demixed PCA (dPCA) components for an example session with monkey N performing normal trials (black), wrist trials (yellow), spring trials (blue), and rubber band trials (green). Solid traces are flexion trials, and dashed traces are extension trials. Components are organized by which behavioral parameter they explain the most neural variance for. (B) Percent of neural variance accounted for by each behavioral parameter. Bars with solid edges are for monkey N, and bars with dashed edges are for monkey W. Error bars indicate standard deviation across sessions. (C) Average muscle activation around peak movement for flexor muscles (top) or extensor muscles (bottom) during trials for each context and target vs the average activation of the first context-dependent dPCA component during the same trials. Each point represents the average activations in a series of trials in one context towards one target. Markers indicate which dataset the sample is from. Correlations are calculated within samples for one target, either flexion (red), or extension (black).

## DISCUSSION

In this study we examined the impact of altering the context of a motor task, either adding an elastic resistance or postural change, while using a BMI for continuous finger control. We found that changes in context increase the error of offline BMI decoder predictions significantly for both kinematics and for muscle activations. This effect was larger for predicting muscle activations than for predicting kinematics. In online trials using a kinematic based BMI, the monkeys were able to quickly adjust for context changes and achieved comparable performance to normal online trials. We tested this in two separate ways, first by adding context changes to the manipulandum during online trials, and second by training two models and swapping between them. In both cases, the monkeys adjusted for the new context as quickly as they adjusted to normal online BMI trials.

The similar online performance in each context, despite large offline mismatch, may be explained by a few possible factors. First would be if normal online trials are performed using a model that already does not capture the relationship between neural activity and intended finger movements well. In a control-systems perspective, online BMI control changes the “plant” from native fingers to the virtual fingers on the screen. There is evidence that as a result of this, neural activity is different during online BMI control (Carmena et al., 2003; Fan et al., 2014; Ganguly et al., 2011; Gilja et al., 2015; Jarosiewicz et al., 2013; Orsborn et al., 2012; Taylor et al., 2002). Additionally, it is unlikely that a linear model like those used here robustly capture the relationship between motor cortex activity and kinematics. Due to the change in neural activity during online trials and inaccuracies in the decoding model, there is likely adjustment required by the monkey to use the BMI online during the normal setup. In this case, performing the online task in a new context is swapping one non-optimal decoder for a new one.

The observed near equivalent online performance could also be observed if the context change does not have a large impact on task-relevant neural activity. Studies into neural plasticity have shown that during a session of online trials, subjects can adjust to decoder perturbations that are within the same intrinsic manifold (Sadtler et al., 2014). We found that individual channel activations change on up to 61.7% of channels that are important for decoding movements (Table I), and this introduces error into model predictions. However, if the perturbations we introduced did not shift activity outside of the intrinsic manifold, then it may have been easy to adjust to the new context. The similar performance with two separate models observed in this study could suggest that our context perturbations did not shift neural activity outside of this manifold. This is supported by our analysis which shows that the intrinsic manifold during normal trials is well aligned with that of trials in other contexts (Figure 6).

The online BMI experiments in this study used a kinematic-based BMI decoder. BMI studies typically predict kinematic variables for applications such as prosthesis control (Hochberg et al., 2012; Wodlinger et al., 2015) and cursor or virtual movement control (Gilja et al., 2015; Hochberg et al., 2006; Young et al., 2019). In the offline predictions using linear models, we found that neither kinematics nor muscle activations could be predicted at the same accuracy in new contexts. While significant, kinematics, specifically flexion velocity, did show a smaller decrease in offline performance between contexts. This may suggest that kinematic variables are more consistent to use as a command signal when kinematics are the final output. However, in FES applications (Ajiboye et al., 2017; Bouton et al., 2016; Nason-Tomaszewski et al., 2022), the large increases in muscle activation required to acquire these off-context targets may be more likely to impact performance. Here the final outputs are stimulation parameters that cause a desired amount of muscle contraction. Importantly, the required stimulation parameters could change with context due to the change in required muscle activation. As a result, even if predictions of position or velocity generalize well to new contexts, the mapping from kinematics to stimulation parameters would no longer be accurate. One solution would be developing a controller to update stimulation parameters to match the decoded command signal, i.e. joint angle or velocity. While FES systems have a lot of variability in performance due to lead placement, subject variability, and muscle fatigue, future work could simulate this system with musculoskeletal models (Chadwick et al., 2009).

Alternatively, generalized FES control could be aided by more accurately decoding intended muscle activations, which can be mapped to stimulation parameters as done by some FES studies (Ethier et al., 2012; Hasse et al., 2022). Non-prehensile finger movements are less studied than arm movements and grasping, partly due to experimental difficulty, with much work coming from only a few data sets (Baker et al., 2009; Schieber, 1991). Although predictions of muscle activations from neural activity for muscles overlapping with those in this study have been done for movements of the wrist (Naufel et al., 2019; Oby et al., 2013) and grasp (Ethier et al., 2012), predicting finger related muscle activations during non-prehensile finger movements has to our knowledge not been attempted yet. Here we found that we could decode muscle activations during this individuated finger task with similar accuracy as decoding kinematics. Predicting muscle activation also led to the poorest offline generalization. The off-context predictions of muscle activation had both a large magnitude change and a lower correlation. The large magnitude change that we observed matches studies of wrist movements where predicting muscle activation also did not generalize well (Naufel et al., 2019). An alternative approach to decoding movements for BMI is to use task-specific features to augment decoder models (Schroeder et al., 2022). Identifying a signal that accounts for the change in muscle activation across contexts would assist in decoder generalization. Here we found that while a large proportion of neural variance was explained by dPCA components that did not change with context, a significant proportion of the neural variance is associated with components that are context-dependent. Visually the context components are shifting the dynamics without changing the overall shape and the shift in neural activity is strongly correlated with muscle activations in new contexts. This could provide a feature for accounting for the gain change while predicting muscle activation or allow models to modulate force or muscle activation while producing the same kinematics. Additionally, some prediction generalization error can be associated with the muscle activations being a higher dimensional variable than kinematics, with evidence that motor cortex can selectively activate or inactivate specific muscles (Schieber et al., 2009). In this study there was no effort to estimate lower dimensional muscle synergies that may be underlying the observed muscle activations, but it is possible that cortical activity would relate more linearly to muscle synergies than to individual muscle activations.

Nonlinear models could also improve predictions of intended muscle activation or kinematics from neural activity. Complex models are becoming more widely used in order to better model the relationship between motor cortex and intended movements (Glaser et al., 2020; Schwemmer et al., 2018; Sussillo et al., 2012; Willett et al., 2021). While a general concern for these nonlinear models is that they will overfit to the training data and not generalize well, given the correct training data they also may be able to identify less obvious trends that will distinguish between contexts and allow for better predictions. For example, Naufel et al. (2019) was able to predict muscle activations in multiple wrist tasks after training an LSTM decoder on data from all of the tasks. This indicates that there may be enough information in our BMI features to distinguish between the different tasks. Our dPCA results indicate that around 24% of neural variance can be accounted for by context specific activity, so it is likely that a neural network would be able to take advantage of that information to make predictions in multiple contexts. More work will be needed to characterize how much training data these nonlinear models need in order to generalize to all the contexts seen at home.

## METHODS

All procedures were approved by the University of Michigan Institutional Animal Care and Use Committee (protocol numbers PRO00010076 and PRO00008138).

### Implants

We implanted two male rhesus macaques (monkey N age 8-9, monkey W age 8-9) with Utah microelectrode arrays (Blackrock Microsystems, Salt Lake City, UT, USA) in the hand area of precentral gyrus, as described previously (Irwin et al., 2017; Nason et al., 2021; Vaskov et al., 2018). Two monkeys were chosen to ensure results are consistent between subjects. Monkey N was implanted with two 64-channel arrays in right hemisphere primary motor cortex and monkey W was also implanted with two 96-channel arrays in right hemisphere primary motor cortex. Channels from both of monkey N’s motor cortex arrays and from monkey W’s lateral motor cortex array were used in this study, for a total of 96 channels from each monkey for analysis. Monkey N was between 511 and 1168 days post-cortical implant and monkey W was between 254 and 411 days post-cortical implant during data collection.

Monkey N was also implanted with chronic bipolar intramuscular electromyography (EMG) recording electrodes (Synapse Biomedical, Inc., Oberlin, OH, USA) in a separate surgery as described previously (Nason et al., 2021). The list of muscles targeted along with their function are included in Table II. Briefly, muscles were accessed via dorsal and ventral incisions on the left forearm and specific muscles were surgically identified. With assistance of intraoperative stimulation, bipolar electrodes were inserted and sutured into the muscle belly near the point of innervation and then tunneled over the elbow and shoulder to an interscapular exit site. The percutaneous electrodes were connected to a 16 channel PermaLoc connector. Monkey N was between 120 and 496 days post-EMG electrode implant for all EMG data collected.

**Table II.**
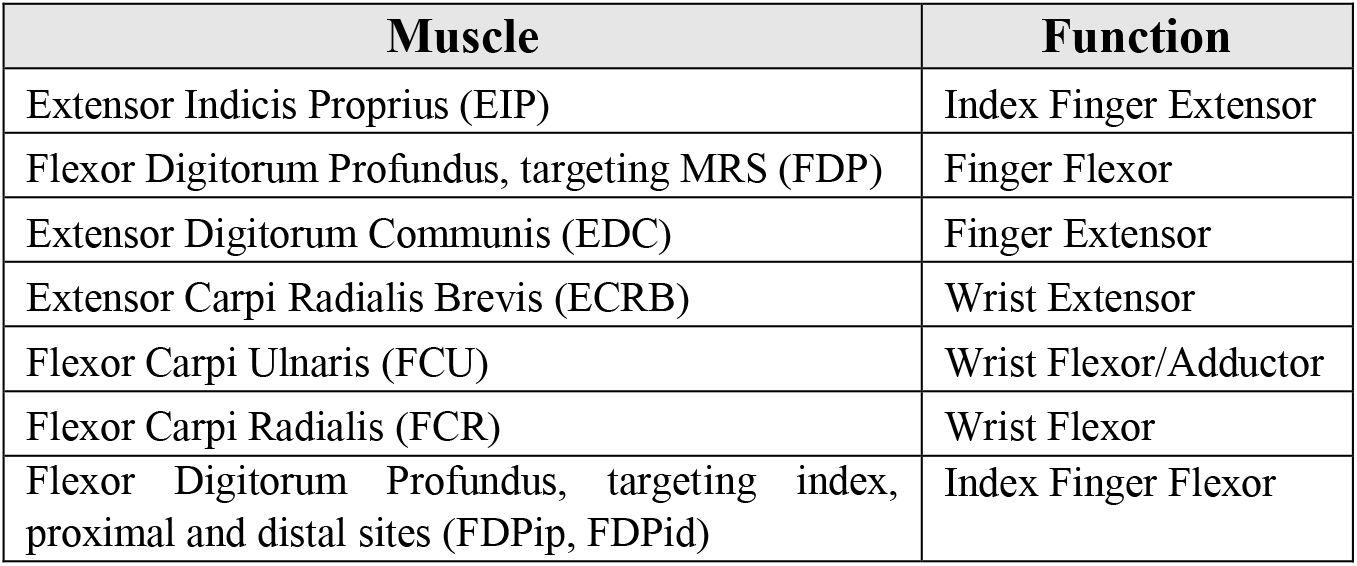
List of muscles targeted during surgery with their associated function.

| Muscle | Function |
| --- | --- |
| Extensor Indicis Proprius (EIP) | Index Finger Extensor |
| Flexor Digitorum Profundus, targeting MRS (FDP) | Finger Flexor |
| Extensor Digitorum Communis (EDC) | Finger Extensor |
| Extensor Carpi Radialis Brevis (ECRB) | Wrist Extensor |
| Flexor Carpi Ulnaris (FCU) | Wrist Flexor/Adductor |
| Flexor Carpi Radialis (FCR) | Wrist Flexor |
| Flexor Digitorum Profundus, targeting index, proximal and distal sites (FDPip, FDPid) | Index Finger Flexor |

### Feature Extraction

Threshold crossing firing rates (TCFR) and spiking band power (SBP) were recorded in real-time during experiments using the Cerebus neural signal processor (Blackrock Microsystems). Threshold crossings were acquired by configuring the Cerebus to threshold each channel at −4.5 times the signal root-mean-square. For each threshold crossing, spike snippets were sent to a computer running xPC Target version 2012b (Mathworks) which saved the channel and time for each threshold crossing. SBP was acquired by configuring the Cerebus to band-pass filter the raw signals to 300-1,000 Hz using the Digital Filter Editor feature in the Central Software Suite version 6.5.4 (Blackrock Microsystems), then sample at two kHz. The filtered 2kHz recording was also sent to the computer running xPC Target, which rectified and summed the samples on each channel received in each 1ms iteration and tracked the quantity of samples received each 1ms. Both the threshold crossings and SBP were saved by xPC synchronized with other real-time experimental information. Artifacts were removed for TCFR by removing threshold crossing times if 20 or more channels had threshold crossings in the same millisecond. For offline analysis, features were binned into non-overlapping bins of length 32ms for online and offline decoding or length of 20ms for calculating tuning and comparing features across trials. SBP is summed for every 1ms in the time bin and then divided by the total number of raw 2kHz samples in the bin. For TCFR the spike counts are summed within a bin and then divided by the bin size to get a threshold crossing rate.

EMG from monkey N’s eight bipolar electrodes was recorded for later offline synchronization. The percutaneous PermaLoc connector was connected to a CerePlex Direct (CPD) via a 64-channel splitter box and CerePlex A (Blackrock Microsystems) which converted the signals to the digital domain with unity gain. The CPD was configured to record 16 channels of raw signal at 10 kHz and for each bipolar pair the electrode implanted further inside the muscle was referenced to the second electrode. These eight bipolar referenced channels are the ones used in analysis while the other eight partner electrodes were still recorded so that the unreferenced signals could be recreated if necessary. To synchronize EMG offline, we used the Sync Pulse functionality in Central to create unique pulses that were recorded by both the Cerebus and CPD and could later be used to align the Cerebus and CPD recordings. For offline analysis, muscle activations are estimated from the 10kHz EMG recording by filtering with a bandpass filter between 100 and 500 Hz and then taking the mean absolute value of the filtered signal during every binning period.

### Experimental Setup

During experiments the monkeys performed a virtual finger task while motor cortex activity and optionally arm muscle activity was recorded as described. Similar to previously described experiments (Irwin et al., 2017; Nason et al., 2021; Vaskov et al., 2018), we used xPC Target to coordinate the experiment in real-time. The xPC Target computer acquired and stored task parameters and neural features in real-time, coordinated target presentation, acquired finger positions from the flex sensors on each finger group (FS-L-0073-103-ST, Spectra Symbol, Salt Lake City, UT, USA), and sent finger positions and target locations to a computer simulating movements of a virtual monkey hand (MusculoSkeletal Modeling Software) (Davoodi et al., 2007). For online experiments, the xPC Target computer also binned threshold crossings and SBP in customizable bin sizes and evaluated the decoder model to predict finger positions in real-time using a ReFIT Kalman filter (see details below).

### Behavioral Task

Monkeys N and W were trained to acquire virtual targets by moving their physical fingers in a manipulandum to control virtual fingers on a screen in front of them. All sessions took place in a shielded chamber with the monkey’s head fixed and arms restrained at their side with elbows bent 90 degrees and hands resting on a table in front of them. The left hand was placed in a manipulandum described previously (Nason et al., 2021), with openings separating the index finger and the middle-ring-small (MRS) finger group. The monkeys were trained to move one finger group independent of another finger group, i.e. 2-degrees-of-freedom (2-DOF), although in some trials they moved both finger groups as one (1-DOF). Each trial began with spherical targets appearing for each active finger group with each target occupying 15% of the full range of motion of the fingers. In the 1-DOF task the target was presented to the index finger.

Target presentation followed a center-out-and-back pattern with every other target presented at a center position, equivalent to 50% on a scale from 0% (full extension) to 100% (full flexion). Additionally, center was presented after any failed trial. The non-center targets were randomly selected from a set of targets. For 2-DOF the targets included any combination of index flexion, rest, or extension, and MRS flexion, rest, or extension, with a randomly chosen magnitude of 20%, 30%, or 40% of the full movement range. The split movements (index flexion with MRS extension or vice versa) did not have a 40% movement magnitude because the monkeys had difficulty splitting the finger groups that far. The 1-DOF movements were also center-out with the fingers flexing or extending either 40% from rest or a randomly chosen magnitude of 20%, 30%, or 40% from rest depending on the session, the former generally being used for tuning analyses and offline comparisons and the latter being used for online experiments. In each trial the monkey had to hold their fingers within the target(s) for 750ms. During online decoding experiments, the same center-out-and-back target presentation order was used but the hold time was reduced to 500ms. In one session used in offline analysis and four online sessions, the hold time was 2ms longer than expected due to a minor bug.

Task context was altered through four potential task alterations. One alteration was the addition of torsional springs to both finger groups (180-degree deflection angle, 0.028in or 0.04in wire diameter, Gardner Spring Inc., Tulsa, OK), referred to as the “spring context”. The second alteration was the rotation of the manipulandum by 23 degrees, referred to as the “wrist context”. A third alteration was introduced by attaching rubber bands from the back of the manipulandum to the door for each finger group, thereby resisting flexion, referred to as the “rubber band context”. A last alteration was addition of torsional springs and the rotation of the manipulandum by 23 degrees at the same time, referred to as the “both context”. Trials performed with one of these alterations are referred to as “off-context” trials, while trials performed without alterations are referred to as “normal” trials. As the index finger alone is much weaker than the MRS finger group, the index finger used a smaller spring when applicable. The added springs increased the force required for full flexion by 9.5N (for MRS) and 3.3N (for index), while the rubber bands increased the force required for full flexion by 16.5N. The rubber band context was only done by monkey N and in a 1-DOF task due to task difficulty.

### Comparison of kinematics and muscle activation between contexts

Three representative sessions for both monkey N and monkey W, one day for each context – spring, wrist, and both – were used to compare kinematics across contexts. During data collection, normal trials and off-context trials were interleaved by alternating context type every 175 to 350 trials in order to control for changes in neural activity over time. During these representative days, there was an average of 1134 normal trials and 1118 off-context trials per day for monkey N and 526 normal trials and 504 off-context trials per day for monkey W. To compute how finger velocity changed between normal trials and off-context trials, the peak velocity of finger movements was found for every trial. For every trial, the recorded finger flexions were down sampled to 20ms and filtered with a second-order Savitzky-Golay FIR filter. Finger velocity was estimated from the down sampled and filtered finger positions and maximum finger speeds were found. The peak movement time was taken at the time of the largest peak in speed after trial start. Trials were then split by context and target direction (flexion vs. extension), and a two-sample *t*-test was used to compare peak speeds and compute a 99% confidence interval, once for flexion targets and again for extension targets. Comparisons were made only between trials to the same target, leaving about 281 trials per group and 129 trials per group for each comparison for monkey N and monkey W respectively.

The same sessions for monkey N used to compare kinematics were also used to compare muscle activations. The recorded EMG was filtered and the mean-absolute value was taken in 20ms bins as described previously. Binned muscle activations were then smoothed with a 100ms Gaussian kernel. One value was obtained for every trial by taking the average muscle activation in a 420ms window around peak movement, including ten bins before peak movement, the bin that included peak movement, and ten bins after peak movement. These muscle activation values were grouped by context and target, then compared with a two-sample t-test.

### Computation of Neural Tuning and Context Modulation

During five representative experiments for each monkey, three that tested the spring context and two that tested the wrist context, we calculated the number of channels that were significantly modulated by any finger movement and the number of channels with a change in activity between normal trials and off-context trials. During these sessions, the monkeys performed the task with all fingers moving together (1-DOF), in a center out task as described, to targets at either plus or minus 40% from center. Trials that were unsuccessful and trials following unsuccessful trials were removed. There was an average of 1072 normal trials and 747 off-context trials for monkey N and 544 normal trials and 387 off-context trials for monkey W were used during these sessions.

Channel tuning and context modulation was calculated with both the SBP features and TCFR features. On each day, features and kinematics were averaged into non-overlapping 20ms bins, data from normal trials and off-context trials were concatenated together, and the SBP and TCFR were each normalized to zero mean and unit standard deviation. An optimal lag was calculated for each channel by maximizing the L2-norm of regression coefficients between a feature and finger position and velocity. Features at that optimal lag were then regressed with finger position and velocity one at a time with an added effect for context following these equations:

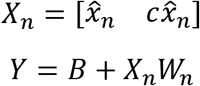

where 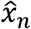 is the T x 1 vector of channel SBP or TCFR for channel n, c is an indicator variable that equals one if that sample was during an off-context trial or zero otherwise, Y is a T x 2 matrix containing finger position and velocity for each bin in T, and W_n_ is the 2 x 2 matrix of trained weights relating channel n’s activity to finger position and velocity. A channel was called tuned if the regression coefficient between the neural feature and either finger position or velocity, i.e. w_1,1_ or w_1,2_, were significantly different from zero, via a t-test on the regression coefficient (p < 0.001). A channel was also called context modulated if either coefficient in the second row of W_n_, which includes the effect of context, was significantly different from zero, also via a t-test (p < 0.001), indicating a different slope relating neural activity and kinematics between normal trials and off-context trials.

To quantify the change in neural activity between contexts as in Figure 6, we used one representative session for monkey N in which trials were done in the normal, spring, wrist, and rubber band contexts. An additional two representative sessions for monkey W were used, one session comparing normal and spring trials and another session comparing normal and wrist trials, both performed with 1-DOF movements and with targets to 40% flexion or extension from rest only. Tuned channels were calculated as previously described using the SBP feature. For every trial, the SBP was binned into 20ms bins and then smoothed with a 100ms Gaussian kernel. Then the average activity in a window spanning 200ms before and 200ms after the bin containing peak movement was calculated for each tuned channel. The trials were then split by context and by target, and the trial SBP values were compared between contexts with a two-sample t-test.

### Offline Predictions

Data from nine sessions with monkey N, three for each context (springs, wrist, and both), were used for offline muscle activation and kinematic predictions, and three sessions for monkey W, one for each context, were used for offline kinematic predictions. During these sessions, both monkeys performed the 2-DOF center-out task. Blocks of normal trials and off-context trials were interleaved by alternating context in order to control for changes in neural activity over time. Trials that were unsuccessful were removed before analysis. There was an average of 803 normal trials and 470 off-context trials for monkey N and 737 normal trials and 329 off-context trials for monkey W. To account for changes in monkey motivation, sessions chosen were those with consistent prediction accuracy between early and late normal trials within a session. These sessions spanned 165 days starting 792 days post-cortical array implant and 120 days post-EMG electrode implant for monkey N, and 63 days starting 285 days post-cortical array implant for monkey W.

In each session, SBP and muscle activations or kinematics were binned into 32ms bins and features were concatenated across trials of the same context. The SBP channels were masked to those with an average TCFR greater than 1Hz across a session and 12 bins of history from each of these channels were used as additional features. Ridge regression relating SBP to muscle activations or kinematics was trained on normal trials and then tested on both normal trials and off-context trials with 10-fold cross validation. To do this, the normal trials were split into 10 folds with an equivalent number of bins in each fold, a model was trained on nine folds, and then tested on the left out fold as well as on data from off-context trials. We used two metrics to evaluate prediction accuracy. First we used the Pearson correlation coefficient between the predicted and measured muscle activations or kinematics to establish how well the predictions are linearly correlated with measurements. The second metric was mean-squared error normalized by the variance of the measured data (MSE). Normalizing by the variance allows for better comparison across test datasets as they may have different variances. In this formulation, the MSE is the fraction of unexplained variance or one minus the variance accounted for or coefficient of determination used in previous studies (Fagg et al., 2009; Naufel et al., 2019). Values greater than one indicate that the predictions are introducing variance compared to the worst possible least-squares predictor, i.e. predicting the mean.

### Online Decoding

We used either a Kalman filter or a ReFIT Kalman filter (Gilja et al., 2012) to predict intended finger movements for all BMI experiments, as done previously (Irwin et al., 2017; Nason et al., 2021; Vaskov et al., 2018). We performed two types of online experiments. In the first experiments, a ReFIT Kalman filter was trained on normal trials and then used during trials with context changes in the manipulandum or without any additions to the manipulandum. To train the model, monkeys first performed at least 300 trials of center-out manipulandum control with 750ms hold time. Using these trials, we trained a position/velocity Kalman filter which the monkeys used online for at least 200 trials, with a 32ms update rate and a 500ms hold time. A ReFIT Kalman filter was then trained, as done previously (Nason et al., 2021), by rotating incorrect velocities during online control with the Kalman filter to be towards the intended target represented in a two-dimensional space, setting finger velocity equal to zero when in the correct target, and then retraining regression coefficient matrices. The ReFIT Kalman filter was used online for blocks of 100 to 200 trials with different context changes applied to the manipulandum, alternating between normal trials and other contexts. Multiple contexts could be tested in one session during these experiments by switching out the context manipulations present in the manipulandum.

In the second set of online experiments, two Kalman filters were trained in one session and then used alternatingly in online control without any changes present in the manipulandum. During these sessions, the monkeys first performed at least 300 trials of center-out manipulandum control, followed by another 300 or more trials of center-out manipulandum control with a context change present. One model was trained using each set of trials. The monkeys then used these models in online control for sets of 100-200 trials, and then the models were alternated. Hold times and update rates were kept consistent between types of experiments and sessions.

### Online Performance Measures

We estimated online performance with acquisition time, time to target, orbiting time, and path efficiency. Acquisition time was measured as the total time from target presentation to the end of the trial minus the hold time, therefore ending with the target being successfully acquired. Time to target was taken as the time from target presentation to the first time where all fingers with targets were in their targets. Orbiting time was then calculated as the time from all fingers first reaching their targets to the end of the trial minus the hold time. Trials where the fingers reached the targets and never left therefore had an orbiting time of 0ms. Failed trials were excluded when comparing online performance between context but not for evaluating the monkeys adaptation within the first five trials. Path efficiency was calculated as the ratio of the shortest distance between the fingers starting position and the target position projected onto a two-dimensional space, to the length of the path traveled by the fingers.

### Dimensionality Reduction

To investigate changes in population neural activity due to changes in context, two sessions of 1-DOF center-out trials with targets of 40% flexion or extension from rest were used for each monkey. For monkey N, both sessions included trials where the task was performed in the normal, spring, rubber band, and wrist contexts. For monkey W, one session included trials in the normal and spring contexts, and the other session included trials in the normal and wrist contexts. SBP was binned into 20ms bins, masked to only include channels with TCFR greater than 1Hz, and then for each trial a time frame 400 ms before to 740 ms after the bin containing peak movement was taken from each trial. The neural activity for trials within a single context was concatenated and averaged across trials with the same context and target forming a N x T x D data structure for each context, where N is the number of channels, T is the number of bins per trial used, and D is the number of targets. Neural data was then concatenated across targets to form an N x (T*D) matrix and then we used PCA to calculate a manifold for each context, keeping the top 16 components for monkey N and eight components for monkey W, which explained 86% percent of variance on average. Principal angles were found between the manifolds following methods used previously (Bjorck & Golub, 1973; Gallego et al., 2018). These principal angles are the minimal angles required to align the manifolds and serve as a measure for how well aligned two manifolds are. As a control, two sets of 50 trials were taken from the normal trials and used to calculate two manifolds in the same way. The principal angles between these manifolds were then calculated. The sampling and angle calculations were repeated 100 times to obtain a control distribution of principal angles.

We also calculated one manifold spanning trials from all contexts tested in one session. This was done using demixed PCA (dPCA) (Kobak et al., 2016). This approach finds a single neural manifold that reduces the dimensionality of the data while maintaining a linear readout that can reconstruct the mean neural activation associated with manually chosen behavioral variables. In this instance, the behavioral parameters chosen were target, i.e. either flexion or extension, and which context the task was done in. MATLAB code for calculating dPCA components was downloaded from http://github.com/machenslab/dPCA. SBP was binned into 20ms bins, masked to include only channels with TCFR greater than 1Hz, and then concatenated into a N x C x D x T x n data structure where N, D, and T follow the same structure as the PCA calculations, n is the number of trials per condition, and C is the number of contexts tested in that session. SBP was averaged over the number of trials, n, to form the peristimulus-time-histograms for each target and context combination, after which dPCA components were calculated. Neural variance of a behavioral parameter was obtained by calculating the variance within the marginalization of neural data based on each behavioral parameter and taking the ratio of the total variance in a marginalization to the total variance in the neural data.

## ACKNOWLEDGEMENTS

We thank Eric Kennedy for animal and experimental support and the University of Michigan Unit for Laboratory Animal Medicine for expert veterinary and surgical support. We appreciate Chris Andrew’s expert statistical assistance and the support of the University of Michigan Biointerfaces Institute. This work was supported by NSF grant 1926576, NSF GRFP under grant 1841052, NIH grant F31HD0998804, NIH grant T32NS007222, NIH grant R01NS105132, the Dan and Better Kahn Foundation Grant AWD011321, University of Michigan Robotics Institute, and A. Alfred Taubman Medical Research Institute.

## DATA AVAILABILITY

Neural, behavioral, EMG, and online BMI performance data will be deposited in an open access repository.

