## Supplemental Tables for "The Impact of Task Context on Predicting Finger Movements in a Brain-Machine Interface"

### Supplementary Results

Supplementary Table I – Average prediction correlation when using a ridge regression model to predict muscle activations or kinematics with the same or different test contexts. Results for monkey N.

| **R** |  | **Monkey N** | | | | | | | | | | | |
| --- | --- | --- | --- | --- | --- | --- | --- | --- | --- | --- | --- | --- | --- |
| **Training Context** | **Test Context** | **FCR** | **FDPid** | **FDPip** | **FDP** | **FCU** | **ECRB** | **EIP** | **EDC** | **Index Position** | **MRS Position** | **Index Velocity** | **MRS Velocity** |
| **Normal** | **Normal** | 0.62 | 0.58 | 0.61 | 0.51 | 0.50 | 0.75 | 0.72 | 0.73 | 0.62 | 0.70 | 0.50 | 0.52 |
| **Wrist** | **Wrist** | 0.62 | 0.54 | 0.60 | 0.49 | 0.51 | 0.68 | 0.66 | 0.66 | 0.55 | 0.71 | 0.51 | 0.52 |
| **Spring** | **Spring** | 0.65 | 0.62 | 0.61 | 0.60 | 0.71 | 0.71 | 0.71 | 0.75 | 0.69 | 0.73 | 0.46 | 0.58 |
| **Wrist+Spring** | **Wrist+Spring** | 0.69 | 0.57 | 0.61 | 0.49 | 0.66 | 0.64 | 0.63 | 0.68 | 0.63 | 0.71 | 0.48 | 0.53 |
| **Normal** | **Wrist** | 0.59 | 0.49 | 0.57 | 0.43 | 0.47 | 0.65 | 0.63 | 0.59 | 0.51 | 0.66 | 0.49 | 0.48 |
| **Normal** | **Spring** | 0.56 | 0.25 | 0.49 | 0.35 | 0.55 | 0.64 | 0.61 | 0.66 | 0.53 | 0.57 | 0.37 | 0.46 |
| **Normal** | **Wrist+Spring** | 0.58 | 0.23 | 0.46 | 0.31 | 0.46 | 0.44 | 0.38 | 0.22 | 0.45 | 0.52 | 0.37 | 0.45 |

| **R** |  | **Monkey W** | | | |
| --- | --- | --- | --- | --- | --- |
| **Training Context** | **Test Context** | **Index Position** | **MRS Position** | **Index Velocity** | **MRS Velocity** |
| **Normal** | **Normal** | 0.43 | 0.40 | 0.49 | 0.32 |
| **Wrist** | **Wrist** | 0.39 | 0.41 | 0.48 | 0.30 |
| **Spring** | **Spring** | 0.59 | 0.34 | 0.51 | 0.44 |
| **Wrist+Spring** | **Wrist+Spring** | 0.43 | 0.44 | 0.44 | 0.39 |
| **Normal** | **Wrist** | 0.37 | 0.38 | 0.49 | 0.28 |
| **Normal** | **Spring** | 0.53 | 0.15 | 0.47 | 0.29 |
| **Normal** | **Wrist+Spring** | 0.32 | 0.29 | 0.40 | 0.33 |

Supplementary Table II – Average prediction correlation when using a ridge regression model to predict kinematics with the same or different test contexts. Results for monkey W.

| **R^2^** |  | **Monkey N** | | | | | | | | | | | |
| --- | --- | --- | --- | --- | --- | --- | --- | --- | --- | --- | --- | --- | --- |
| **Training Context** | **Test Context** | **FCR** | **FDPid** | **FDPip** | **FDP** | **FCU** | **ECRB** | **EIP** | **EDC** | **Index Position** | **MRS Position** | **Index Velocity** | **MRS Velocity** |
| **Normal** | **Normal** | 0.39 | 0.33 | 0.38 | 0.26 | 0.26 | 0.56 | 0.53 | 0.53 | 0.39 | 0.50 | 0.25 | 0.27 |
| **Wrist** | **Wrist** | 0.38 | 0.29 | 0.35 | 0.23 | 0.26 | 0.47 | 0.43 | 0.43 | 0.31 | 0.50 | 0.26 | 0.27 |
| **Spring** | **Spring** | 0.43 | 0.39 | 0.38 | 0.36 | 0.50 | 0.50 | 0.51 | 0.56 | 0.47 | 0.54 | 0.21 | 0.33 |
| **Wrist+Spring** | **Wrist+Spring** | 0.47 | 0.33 | 0.38 | 0.24 | 0.43 | 0.40 | 0.39 | 0.46 | 0.40 | 0.51 | 0.22 | 0.28 |
| **Normal** | **Wrist** | 0.30 | 0.13 | 0.22 | 0.05 | 0.12 | 0.17 | 0.16 | 0.00 | 0.24 | 0.32 | 0.24 | 0.23 |
| **Normal** | **Spring** | 0.11 | -0.89 | -0.26 | -1.78 | -0.18 | -0.14 | -0.26 | 0.32 | 0.08 | 0.06 | 0.10 | 0.18 |
| **Normal** | **Wrist+Spring** | -0.09 | -1.96 | -1.35 | -2.63 | -1.06 | -2.61 | -1.82 | -0.60 | -0.45 | 0.08 | 0.12 | 0.20 |

Supplementary Table III – Average prediction coefficient of determination when using a ridge regression model to predict muscle activations or kinematics with the same or different test contexts. Results for monkey N.

| **R^2^** |  | **Monkey W** | | | |
| --- | --- | --- | --- | --- | --- |
| **Training Context** | **Test Context** | **Index Position** | **MRS Position** | **Index Velocity** | **MRS Velocity** |
| **Normal** | **Normal** | 0.19 | 0.16 | 0.24 | 0.10 |
| **Wrist** | **Wrist** | 0.15 | 0.16 | 0.22 | 0.08 |
| **Spring** | **Spring** | 0.35 | 0.11 | 0.25 | 0.19 |
| **Wrist+Spring** | **Wrist+Spring** | 0.18 | 0.19 | 0.18 | 0.14 |
| **Normal** | **Wrist** | 0.06 | 0.11 | 0.23 | 0.08 |
| **Normal** | **Spring** | -1.46 | -0.16 | 0.09 | 0.08 |
| **Normal** | **Wrist+Spring** | -0.67 | -0.09 | 0.00 | 0.10 |

Supplementary Table IV – Average prediction coefficient of determination when using a ridge regression model to predict kinematics with the same or different test contexts. Results for monkey W.
